# NF-κB-c-REL impairment drives human stem cells into the oligodendroglial fate

**DOI:** 10.1101/664060

**Authors:** Lucia M Ruiz-Perera, Johannes FW Greiner, Christian Kaltschmidt, Barbara Kaltschmidt

## Abstract

Molecular mechanisms underlying fate decisions of human neural stem cells (NSCs) between neurogenesis and gliogenesis are critical during neuronal development and progression of neurodegenerative diseases. Despite its crucial role in murine nervous system development, the potential role of the transcription factor nuclear factor kappa-light-chain-enhancer of activated B-cells (NF-κB) in fate shifts of human stem cells is poorly understood.

Facing this challenge, we demonstrate here that NF-κB-c-REL drives glutamatergic differentiation of adult human stem cells, while its impairment results in a shift towards the oligodendroglial fate. We particularly observed an opposing balance switch from NF-κB-RELB/p52 to NF-κB-c-REL during early neuronal differentiation of NSCs originating from neural crest-derived stem cells. Exposure of differentiating human NSCs to the c-REL inhibiting approved drug pentoxifylline (PTXF) resulted in elevated levels of cell death and significantly decreased amounts of NF200^+^/VGLUT2^+^ neurons. PTXF-mediated inhibition of c-REL further drove human NSCs into the oligodendrocyte fate, as demonstrated by a complete switch to OLIG2^+^/O4^+^ oligodendrocytes, which also showed PDGFRα, NG2 and MBP transcripts.

In summary, we present here a novel human cellular model of neuronal differentiation with an essential role of NF-κB-c-REL in fate choice between neurogenesis and oligodendrogenesis potentially relevant for multiple sclerosis and schizophrenia.

## Introduction

Development, function and regeneration of the human brain crucially depend on the generation of neurons, astrocytes and oligodendrocytes by neural stem cells (NSCs) (Ma *et al.*, 2009). Differentiation of NSCs results in the isometric population of neurons and glia observed in the human brain (Azevedo *et al.*, 2009) with particular proportions between distinct glial cell populations. For instance, in the cerebral cortex astrocytes corresponded to 20%, microglia to 5% and oligodendrocytes to even 75% of all glial cells (Pelvig *et al.*, 2008). The mechanisms underlying brain development and composition during adulthood crucially depend on fate decisions of NSCs, which are in turn also directly linked to progression of neurodegenerative diseases. For instance, Alzheimer’s disease was very recently correlated to the disability of NSCs to undergo neuronal differentiation (Ribeiro *et al.*, 2019). In terms of multiple sclerosis, differentiation of NSCs into oligodendrocytes is suggested to contribute to myelin repair in addition to the role of resident oligodendroglial precursor cells (Miron *et al.*, 2011). In addition, cognitive ageing decline is correlated with a decrease in oligodendrocytes and neurons but not astrocytes (Pelvig *et al.*, 2008), further indicating the importance to understand the molecular mechanisms of lineage choices of NSCs and their role in aging and disease. The transcription factor nuclear factor kappa-light-chain-enhancer of activated B-cells (NF-κB) is involved in diverse cellular processes like inflammation or cell death, but also plays a particular critical role in the mammalian brain during development and adulthood. Especially the NF-κB subunit p65 was observed to have a predominant activity in the adult mouse brain, while its activity was directly linked to synaptic function, transmission and plasticity (Kaltschmidt *et al.*, 1993, 1995; Kaltschmidt *et al.*, 1997; Imielski *et al.*, 2012) as well as axonal outgrowth and polarization (Sanchez-Ponce *et al.*, 2008; Imielski *et al.*, 2012). NF-κB-p65 was also activated during neural differentiation of mouse NSCs, while its inhibition promoted self-renewal (Zhang *et al.*, 2012). However, little is known about the role of NF-κB in the regulation of human NSCs during neuronal differentiation and their respective lineage choices between neurogenesis and gliogenesis.

In the present study, we took advantage of a human neural stem cell model, which we very recently used to investigate the NF-κB p65-dependent neuroprotection in human neurons (Ruiz-Perera *et al.*, 2018). We isolated neural crest-derived stem cells (NCSCs) from the adult human nasal cavity (Greiner *et al.*, 2011; Hauser *et al.*, 2012) and differentiated them into NSCs, and further into functional mature glutamatergic neurons according to our previous studies (Ruiz-Perera *et al.*, 2018). A comparable mouse model system was previously used to study fate shifts of NSCs originating from NCSCs (Weber *et al.*, 2015). Applying this ideal model to study human neuronal development and maturation, we demonstrate here that glutamatergic differentiation of human NCSC-derived NSCs is predominantly driven by NF-κB-c-REL. Inhibition of c-REL by pentoxifylline (PTXF), a approved drug shown to deviate the immune response in multiple sclerosis patients (Rieckmann *et al.*, 1996), induced a direct shift from the neuronal towards the oligodendrocyte fate, evidenced by the increase of oligodendrocyte specific markers OLIG2, O4 and MBP. These findings demonstrate that c-REL is required for human neuronal fate commitment, particularly concerning the neuronal and oligodendroglial fate and suggest an indispensable role during demyelinating diseases like multiple sclerosis.

## Materials and Methods

### Isolation and Cultivation of human NCSCs

NCSCs were isolated from adult human inferior turbinate tissue obtained by biopsy during routine surgery after informed consent according to local and international guidelines and cultivated as described previously (Greiner *et al.*, 2011; Hauser *et al.*, 2012). All experimental procedures were ethically approved by the ethics board of the medical faculty of the University of Münster (No. 2012–015-f-S).

### Neuronal differentiation of human NCSCs from NSCs to glutamatergic neurons

For guided differentiation of NCSCs to NSCs and further to glutamatergic neurons, NCSCs cultivated as described above were exposed to neuronal induction medium as previously described (Ruiz-Perera *et al.*, 2018). For detailed procedure see supplementary material and Fig. 1A.

**Figure 1.**
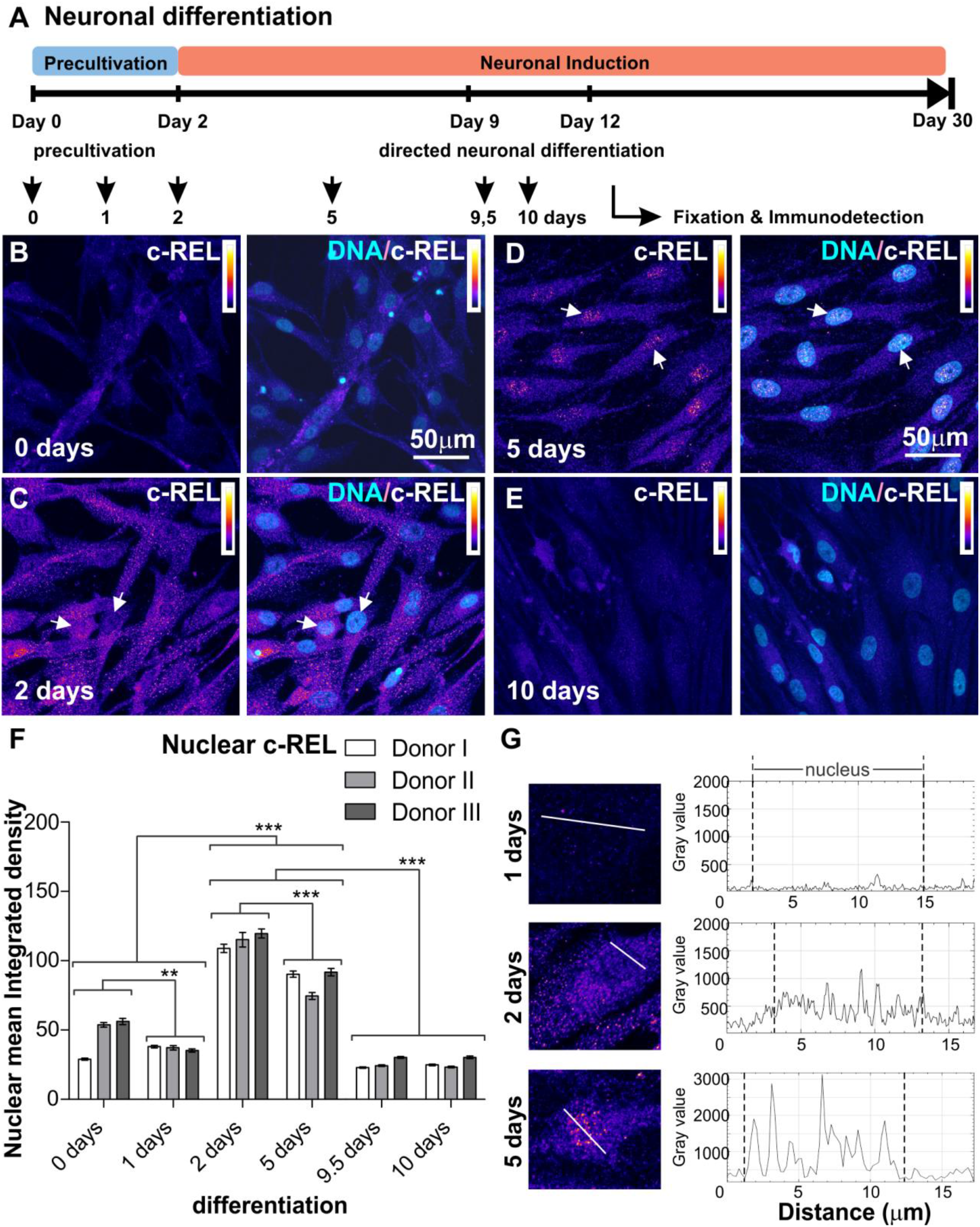
Immunocytochemical analysis of c-REL. A) Schematic overview of neuronal differentiation procedure. Adult human NCSC-derived NSCs were differentiated into glutamatergic neurons, samples were taken atdifferent time points to analyze the NF-κB subunit composition during early differentiation A-E) NCSC-derived NSCs labeled against c-REL after 0, 2, 5, and 10 days of glutamatergic differentiation respectively. Each panel shows on the left c-REL and co-localization with DNA on the right. Intensity scale indicates white as highest intensity level and black as lowest. Arrows depict c-REL nuclear activation. F) Quantification of immunocytochemical analyses showing nuclear mean integrated density for c-REL during early differentiation (n=3, mean ± SEM). Normality of the data was refuted using Shapiro-Wilk normality test. Non-parametric Kruskal-Wallis (***p≤0.001) and Bonferroni corrected post-test (***p<0.001) revealed significantly increased nuclear translocation of NF-κB-c-REL at day 2 and day 5. G) Fluorescence intensity profiles measured at three different time points (1, 2 and 5 days of differentiation), for different cells following transects as shown, in order to clearly reveal the difference between the nuclear and cytoplasmic fluorescence.

### Pentoxifylline treatment

Inhibition of c-REL-activity via PTXF-treatment was performed by adding 500 μg / ml PTXF (Grinberg-Bleyer *et al.*, 2017) to the media in parallel to the neuronal differentiation procedure as described above. PTXF was refreshed every 1-2 days for 30 days, while differentiating NSCs not exposed to PTXF were used as control.

### Immunocytochemistry

Differentiated NCSCs were fixed in phosphate-buffered 4% paraformaldehyde (pH 7.4) for 15 minutes at room temperature following immunocytochemical staining procedure as previously described (Ruiz-Perera *et al.*, 2018). For detailed procedure as well as used primary and secondary antibodies see supplementary material. Fluorescence imaging was performed using confocal laser scanning microscopy (LSM 780; Carl Zeiss, Jena, Germany) and analyzed using ZEN software (Carl Zeiss) or ImageJ.

### Quantitative Polymerase chain reaction (qPCR)

Total RNA was isolated using NucleoSpin RNA kit (Macherey-Nagel, Düren, Germany) according to the manufacture’s guidelines. Quality and concentration of RNA were assessed via Nanodrop ultraviolet spectrophotometry followed by cDNA synthesis using First Strand cDNA Synthesis Kit (Fermentas, Thermofisher Scientific, Waltham, MA, USA). Quantitative PCR were performed in triplicates using PerfeCTa SYBR® Green SuperMix (Quantabio, Beverly, MA, USA) according to the manufacturer’s guidelines and assayed with a Rotor Gene 6000 (Qiagen, Venlo, Netherlands). Primers are listed in Supplementary table S1.

### Image analyses and quantification

For analysis of nuclear NF-κB protein, quantification of indirect immunofluorescence was performed for a minimum of 3 different donors by analyzing 6-12 pictures per time point and donor. Mean nuclear integrated density was measured by defining the region of interest with the nuclear counterstaining using ImageJ. For analysis of neuronal survival, the amount of nonviable neurons recognized by nuclear condensation and/or fragmented chromatin was counted and death rate was calculated as previously described (Ruiz-Perera *et al.*, 2018).

### Statistical analyses

Statistical analysis was performed using Past3 and GraphPad Prism 5 (GraphPad Software, La Jolla, CA). Normality of the data sets was refuted after analysis using Shapiro-Wilk normality test. Homogeneity of variance was tested using Levene’s test and non-parametric Kruskal-Wallis test was applied to compare medians between the different data sets for the different donors (***p value<0.001). Non-parametric Mann-Whitney test was used to compare two pair of groups (***p≤0.001). Tukey’s test or Bonferroni corrected post-test served to identify the significance of the differences between the groups, by comparing the medians or means respectively (*p≤0.05, **p≤0.01,***p≤0.001).

### Data availability

The authors confirm that the data supporting the findings of this study are available within the article and its supplementary material.

## Results

### NF-κB-c-REL orchestrates the early stages of glutamatergic differentiation in NCSC-derived neural stem cells

Neural stem cells originating from neural crest-derived stem cells were exposed to neuronal differentiation conditions for up to ten days according to our previously described protocol (Ruiz-Perera *et al.*, 2018; see Fig. 1A for overview). Activities of NF-κB subunits p65, RELB, c-REL, p50 and p52 (Fig. S1A) were characterized by assessing their nuclear localization (Fig. S1B) via immunocytochemistry. We observed low levels of nuclear p65 protein, which even revealed a further significant decrease during early neuronal differentiation for up to ten days (Fig. S2). A slight transient increase in nuclear RELB at day one of differentiation was accompanied by elevated protein levels of nuclear p52, which were both found to be significantly reduced after 24 h and during further neuronal differentiation of NSCs (Fig. S3 and S4). While p50 protein levels showed no relevant differences during neuronal differentiation (Fig. S5), levels of IκBα protein were elevated from 2 to 5 days of neuronal differentiation, indicating inactivation of RELB/p52 dimers in an IκBα-dependent manner (Fig. S6). Notably, a strong and highly significant increase of nuclear c-REL protein was observable from day 2 to day 5 of neuronal differentiation (Fig. 1B-G). In summary, our findings indicate an opposing balance switch from RELB/p52 towards c-REL-activity to orchestrate the early phase of neuronal differentiation of human NSCs (Fig. 2A).

**Figure 2.**
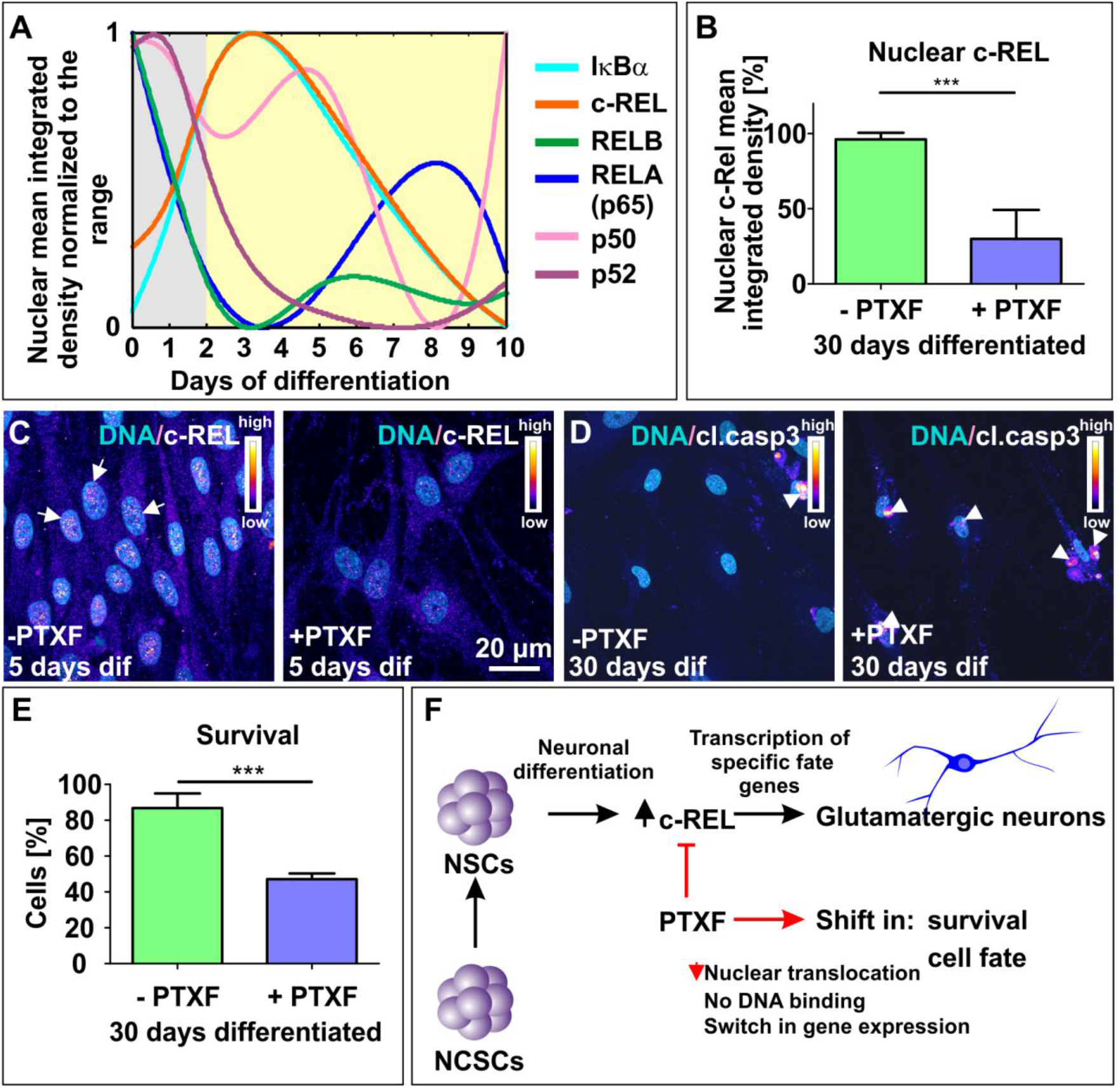
NF-κB subunit composition in early gluatamatergic differentiation and c-REL potential function. A) NF-κB subunit distribution in early stages of glutamatergic differentiation. Nuclear mean integrated density illustrated as the median normalized to the range in a scale from 0 to 1, each NF-κB subunit curve as indicated by colours. B) Nuclear mean integrated density quantified for c-REL immunocytochemistry (n=3, mean ± SEM). Non-parametric Mann Whitney test confirmed a significant decrease in c-REL nuclear translocation after 5 days of differentiation with PTXF (***p≤0.0001), a known c-REL inhibitor (Grinberg-Bleyer *et al.*, 2017). C) 5 days differentiated NCSC-derived NSCs with and without PTXF, labeled against c-REL shown with intensity scale and merged with DNA. Arrows depict c-REL nuclear activation. D) NCSC-derived NSCs differentiated for 30 days without and with PTXF, labeled against clived-caspase-3 (cl.casp-3, intensity scale) and merged with nuclear counterstaining. E) Survival quantified for differentiated NCSC-derived NSCs after 30 days in the presence or absence of pentoxifylline (PTXF; n=3, mean ± SEM). Non-parametric Mann Whitney test confirmed a significant decrease in survival during PTXF-treatment (***p≤0.001), compared to untreated neurons. F) Diagram showing neuronal differentiation mechanism and PTXF mechanism of action. NCSCs differentiate into neural stem cells.These NSCs differentiating into neurons undergo an increase in nuclear NF-κB-c-REL, which further induces differentiation in the glutamatergic neuronal fate. The treatment using PTXF during the early neuronal differentiation diminishes c-REL activation leading to a decrease in the gene expression towards neuronal fate resulting in a shift in the cell fate or otherwise cell death.

### NF-κB-c-REL is crucial for cell survival during glutamatergic differentiation of adult human stem cells

To further determine the role of NF-κB-c-REL during differentiation of NCSC-derived NSCs into the glutamatergic neurons, we exposed differentiating NSCs to c-REL-activity inhibitor and approved drug pentoxifylline (PTXF; Grinberg-Bleyer *et al.*, 2017). We confirmed inhibition of c-REL-activity by observing significantly decreased amounts of nuclear c-REL protein in PTXF-treated differentiating NSCs (day 5 of differentiation) in comparison to non-treated controls (Fig. 2B-C). Notably, PTXF treatment resulted in increased amounts of cleaved-caspase3 positive neurons after prolonged differentiation of NSCs into glutamatergic neurons for 30 days compared to control (Fig. 2D). Quantification of neuronally differentiated cells showing nuclear condensation and/or fragmented chromatin confirmed a significant decrease in the survival of the PTXF-treated differentiated glutamatergic neurons (47.14 ± 3.17% survival, Fig. 2E) compared to untreated neurons (86.83 ± 8.16% survival, Fig. 2E). These findings demonstrated that NF-κB-c-REL has a central role in cell survival during differentiation of human stem cells into the neuronal fate and raised the hypothesis whether this may be accompanied by a shift in the cell fate (Fig. 2F).

### c-REL inhibition by pentoxifylline induces a shift into the oligodendrocyte fate

We investigated potential c-REL-dependent cell fate switches in differentiating NSCs exposed to PTXF by analyzing mature neuronal and glial markers as schematically depicted in Figure 3A. Here, treatment of differentiating NSCs with PTXF led to significantly reduced amounts of NF200^+^ (41,44% ± 6.31%) / VGLUT2^+^ (47.97% ± 2.72%) glutamatergic neurons in comparison to control (84,68% ± 7,70% NF200^+^ / 85.37% ± 5.34% VGLUT2^+^ neurons) (Fig. 3B-C). Interestingly, PTXF-treated differentiating NSCs were completely negative for glial fibrillary acidic protein (GFAP), suggesting the absence of astrocytes (Fig 3D). On the contrary, PTXF-treated NSCs underwent differentiation into the oligondendrocyte lineage, as demonstrated by a complete and highly significant shift towards OLIG2^+^ cells (86,67% ± 6,67%), which were not detectable in the non-treated control (Fig. 3E). Significantly increased amounts of O4^+^ cells after PTXF-treatment (100%) compared to untreated control (Fig. 3F) were accompanied by the presence of Platelet derived growth factor receptor alpha (PDGFRα), transmembrane proteoglycan nerve-glia antigen 2 (NG2) and Myelin basic protein (MBP) at the mRNA level, which were totally absent in untreated neurons (Fig. 4A-C). The observed cell fate shift was further confirmed by a major change in cellular morphology of PTXF-treated cells which acquired a polygonal-like shape with an extended cell surface as well as a reduction of the cellular processes in comparison to control (Fig 3B-E). In addition, we observed a slightly increased amount of Nestin^+^ / p75^+^ cells after PTXF-treatment, suggesting a rather undifferentiated state in comparison to NSCs-derived glutamatergic neurons (Fig. S7A-B). PTXF-treated and untreated NSCs were further found to be negative for alpha smooth muscle actin (αSMA), indicating the absence of mesodermal cells and a respective fate switch towards the mesodermal lineage (Fig. S7C). In summary, our findings demonstrate a NF-κB-c-REL-mediated fate shift of adult human stem cells from glutamatergic neurons towards oligodendrocytes (Fig. 4D).

**Figure 3.**
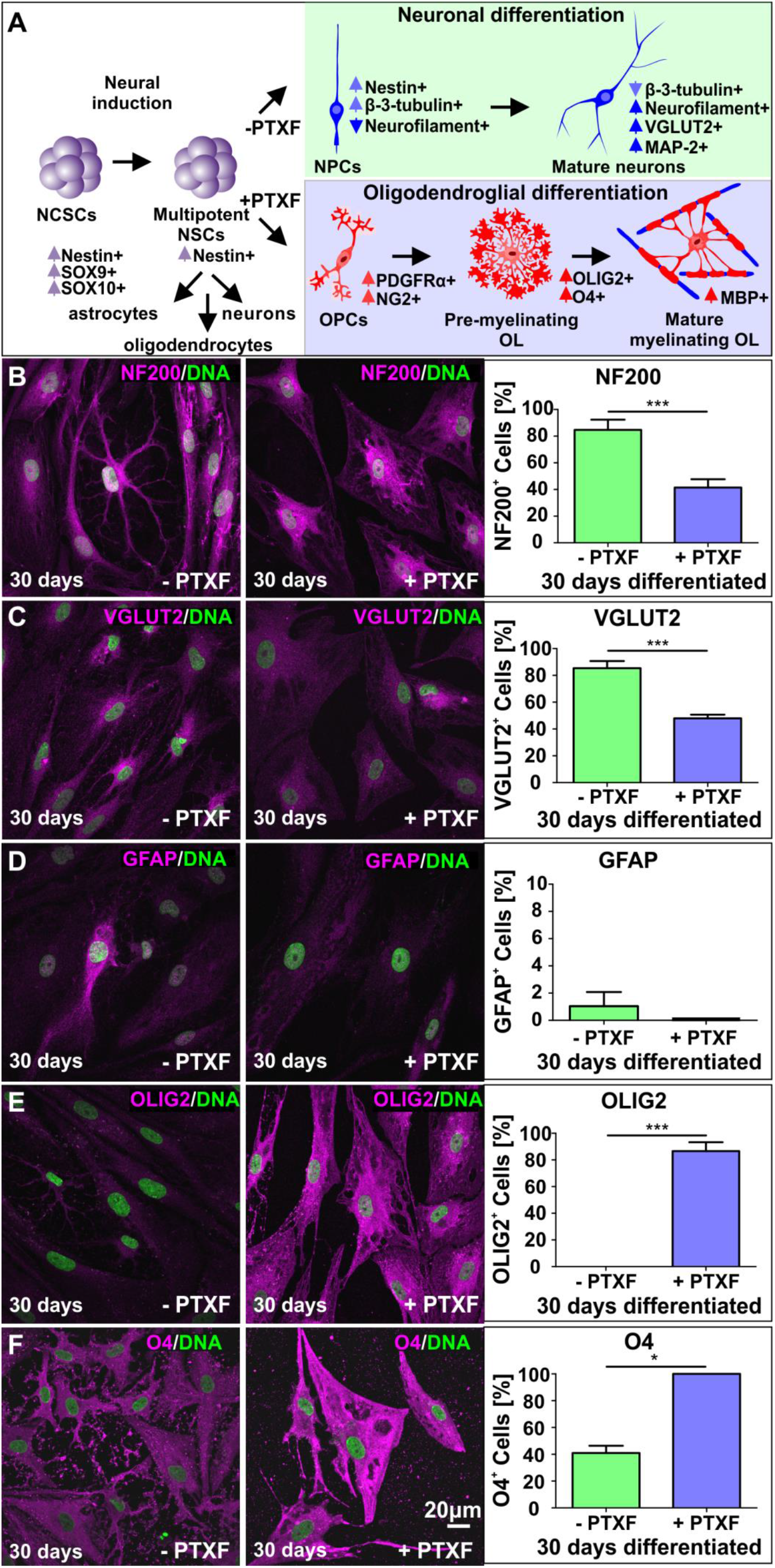
Immunocytochemistry assays showing neuronal and glial markers after 30 days of glutamatergic differentation in presence or absence of pentoxifylline and respective quantifications. A) Schematic overview showing changes in transcription expression during specification of NCSCs into multipotent NSCs that further differentiate into astrocytes, oligodendrocytes and neurons. Differentiation in the absence or presence of PTXF directed the cell fate into the neuronal (light green) or oligodendroglial fate (light blue), further analyzed by the expression of particular markers as indicated (modified from Miron *et al.*, 2011; Hauser *et al.*, 2012). B) Neuronal differentiated NCSC-derived NSCs labeled against Neurofilament 200 (NF200, high molecular weight subunit), and quantification showing percentage of NF200^+^ neurons. Non-parametric Kruskal-Wallis test (***p<0.0005) revealed a significant decrease of NF200^+^ differentiated-neurons treated with PTXF (41,44% ± 6.31%), compared to the untreated neurons (84,68% ± 7,70%). C) Neuronal differentiated NCSC-derived NSCs labeled against VGLUT2, and quantification showing the percentage of VGLUT2^+^ neurons. Non-parametric Kruskal-Wallis test (***p<0.0005) confirmed a significant decrease of VGLUT2^+^ neurons in the differentiated cells treated with PTXF (47,97% ± 2,72%), in comparison to the untreated neurons (85,37% ± 5,34%). D) Differentiated NCSC-derived NSCs labeled against GFAP (astrocyte marker) and respective quantification. Percentage of GFAP^+^ cells indicates a very low number of astrocytes in the untreated-neurons (1,04% ± 1.04%) and complete absence in PTXF-treated differentiated NCSC-derived NSCs (0%, no significant difference). E) Neuronal differentiated NCSC-derived NSCs labeled against OLIG2 (early oligodendrocyte marker) and quantification of OLIG2^+^ cells. Which showed null percentage of positive cells in untreated neurons (0%), while in differentiated-treated-NSCs almost every cell was OLIG2+ (86,67% ± 6,67%). Non-parametric Kruskal-Wallis test (***p<0.001) confirmed significant increase of OLIG2^+^ cells upon PTXF treatment, indicating a shift into the oligodendrocyte fate. F) Neuronal differentiated NCSC-derived NSCs labeled against O4 (immature oligodendrocyte marker) and quantification of O4^+^ cells, showing an increase of O4^+^ cells in PTXF-treated neuronal differentiated NCSC-derived NSCs (100%), confirmed by statistical non-parametric Kruskal-Wallis test (*p<0.05) compared to untreated neurons (40,97% ± 5.42%). PTXF: pentoxifylline, NCSCs: neural crest-derived stem cells, NSCs: neural stem cells, OPCs: Oligodendrocyte precursor cells, OL: oligodendrocytes,

**Figure 4.**
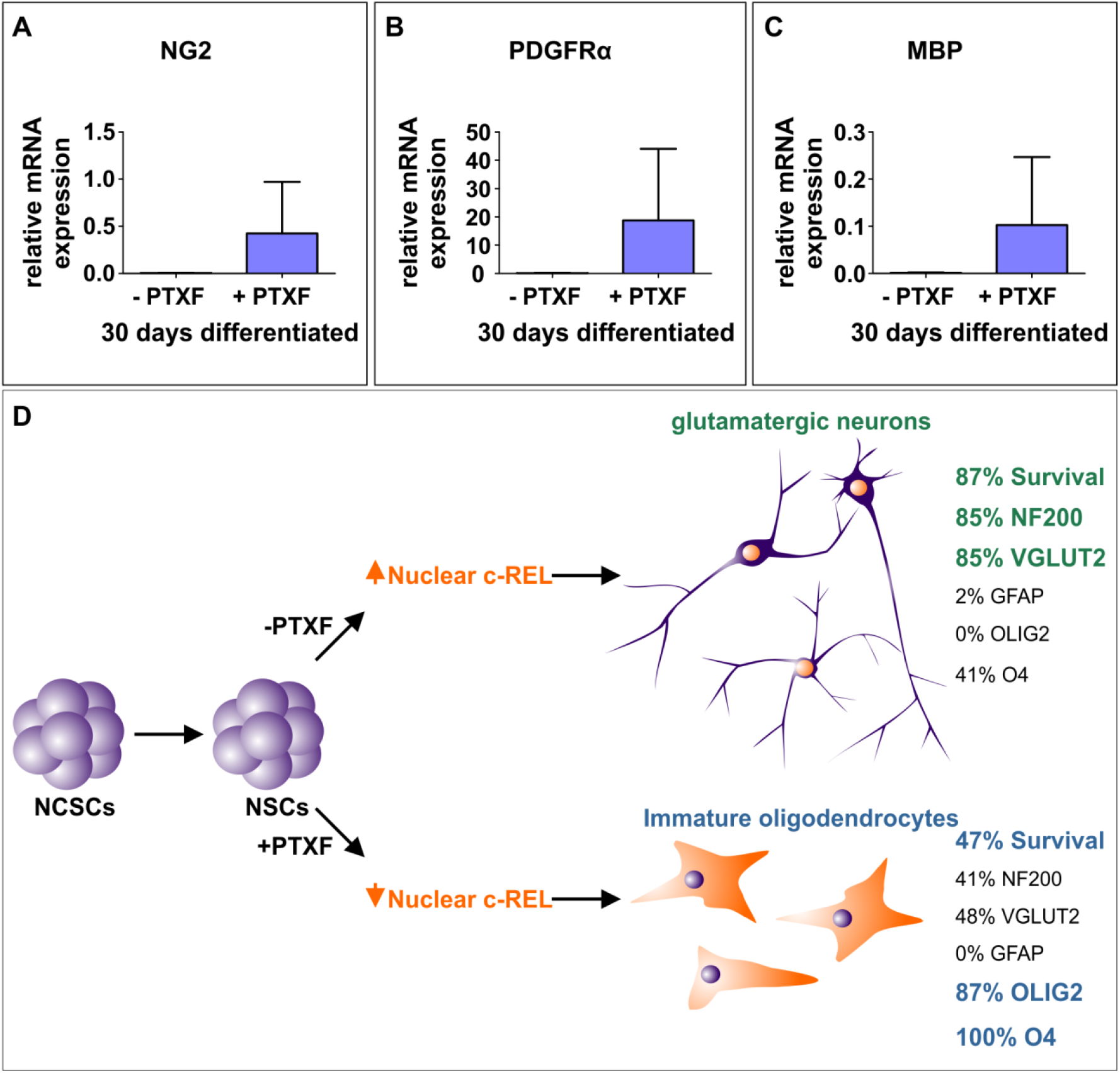
c-REL early inhibition induces a cell fate shift towards the oligodendrocyte phenotype. A-C) Real time polymerase chain reaction showing relative mRNA levels of Oligodendrocyte markers NG2, PDGFRA and MBP respectively compared to the mean value of the reference genes GAPDH and RPLP0. mRNA for these early (NG2, PDGFRa) and mature oligodendrocyte markers (MBP) are only present in the differentiated NCSC-derived NSCs treated with PTXF after their differentiation, indicating a shift into the oligodendrocyte lineage (n=2, mean ± SD). D) Summarizing scheme showing outstanding results obtained. NCSCs are specified into NSCs, and upon neuronal differentiation these NSCs differentiate into glutamatergic neurons. Cells undergo a strong increase of c-REL activation during early time points in the absence of PTXF, which further induces the expression of a specific gene program towards the neuronal fate. This was confirmed by the increased expression of mature neuronal markers NF200 and VGLUT2, and a clear decrease in O4, as well as almost no GFAP and OLIG2 accompanied by a strong survival. When differentiating NSCs were treated in parallel with PTXF, the increase of nuclear c-REL was completely abolished, leading to a shift in the cell fate into the oligodendrocyte lineage, which was confirmed by a strong increase in oligodendrocyte markers OLIG2 and O4 and a clear decrease in the neuronal markers NF200 and VGLUT2. This was also accompanied by a strong reduction in cell survival. PTXF: pentoxifylline, NCSCs: neural crest-derived stem cells.

## Discussion

Our findings demonstrate for the first time that glutamatergic differentiation of human NCSC-derived NSCs is driven by NF-κB-c-REL, while c-REL impairment drives a direct shift into the oligodendrocyte fate. We particularly observed an opposing balance switch from RELB/p52 to c-REL in an early phase of glutamatergic differentiation, orchestrating fate decisions towards the neuronal fate. While the observed activity switch from RELB/p52 to c-REL was mediated in an IκBα-dependent manner, RELA and p50 showed no activity during this process. Accordingly, activity of NF-κB-p65 was implicated in maintenance of pluripotency and decreased upon differentiation of human embryonic stem cells (hESCs) (Armstrong *et al.*, 2006; Takase *et al.*, 2013). In addition, a similar decrease in NF-κB-p65 allowed differentiation of hESC-derived neural progenitor cells during *in vitro* neurogenesis (FitzPatrick *et al.*, 2018). Knockdown of p65 in the initial differentiation phase of hESCs likewise resulted in the upregulation of endodermal and mesodermal markers and mesenchymal specification (Deng *et al.*, 2016). In contrast, p65-activation was reported to be increased upon differentiation in mouse embryonic stem cells (Kim *et al.*, 2008; Lüningschrör *et al.*, 2012). Despite this well-characterized role of NF-κB-p65 in defining the undifferentiated state in human and mouse embryonic stem cells, p65 seems not to directly mediate cell fate decisions in human stem cells. Accordingly, our present findings define a novel role of NF-κB-c-REL in mediating differentiation, neuroprotection and fate shifts of adult human stem cells. In terms of neuroprotection, we observed a significantly reduced cell survival during differentiation of hNSCs in the presence of the c-REL inhibitor PTXF. These results are in accordance with previously described anti-apoptotic effects of c-REL in neurons (Pizzi *et al.*, 2002) and the role of NF-κB as a regulator of cell proliferation (Widera *et al.*, 2006).

Next to the loss of neuroprotection, PTXF-dependent inhibition of c-Rel-activity directly induced a switch from neurogenesis to gliogenesis *in vitro*, particularly into the oligodendrocyte fate. Inhibition of c-REL allowed hNSCs to differentiate into a heterogeneous cell population containing PDGFRα^+^/NG2^+^-oligendendrocyte precursor cells, immature OLIG2^+^/O4^+^-pre-myelinating oligodendrocytes and mature myelinating oligodendrocytes expressing MBP, according to the classification of oligodendrocyte-specific markers by Miron and coworkers (Miron *et al.*, 2011). Interestingly, a similar shift from neurogenesis to gliogenesis was observed in the central nervous system (CNS) by downregulation of pro-neuronal genes required for neuronal differentiation (Tomita *et al.*, 2000). Thus, the c-REL-mediated neuronal differentiation of hNSCs as well as the switch towards gliogenesis upon c-REL-inhibition observed here might be dependent on its direct regulation of similar pro-neuronal genes. Despite the strong fate shift towards gliogenesis, hNSCs did not show any signs of differentiation into astrocytes upon inhibition of c-REL. Accordingly, a strong correlation between the number of oligodendrocytes and neurons was observed in the human brain throughout aging, while the amount of astrocytes remained unchanged (Pelvig *et al.*, 2008). These observations are contrary to those made in the mouse brain and emphasize the need to address developmental research questions using human model systems (Valerio-Gomes *et al.*, 2018).

Our present findings suggest that c-REL might be indispensable *in vivo* and play a particular role in neurodegenerative diseases like multiple sclerosis and schizophrenia. Interestingly, in schizophrenia a 28% reduction in the number oligodendrocytes was observed in human patients (Hof *et al.*, 2003). PTXF-treatment was shown to deviate the immune response in multiple sclerosis patients by decreasing the levels of pro-inflammatory cytokines like TNFα and interleukin-2 (Rieckmann *et al.*, 1996). Next to this commonly known mode of action, our present observations suggest that PTXF may directly modify the cell populations within the CNS via c-REL-inhibition *in vivo*. Emphasizing the role of fate shifts in multiple sclerosis, fibrinogen was recently observed to enter the brain via a damaged blood-brain-barrier and shift the differentiation of OPCs from oligodendrocytes towards astrocytes, further decreasing remyelination (Petersen *et al.*, 2017; Nave and Ehrenreich, 2019).

In summary, we demonstrate here an essential role of NF-κB-c-REL in the choice of human NSCs between neuronal and oligodendroglial fate, suggesting an indispensable role during neural differentiation *in vivo* and demyelinating diseases like multiple sclerosis.

## Acknowledgments

The excellent technical help of Angela Kralemann-Köhler is gratefully acknowledged.

