## Supplementary Material for "NF-κB-c-REL impairment drives human stem cells into the oligodendroglial fate"

#### Supplementary figures and tables

**Figure S1**

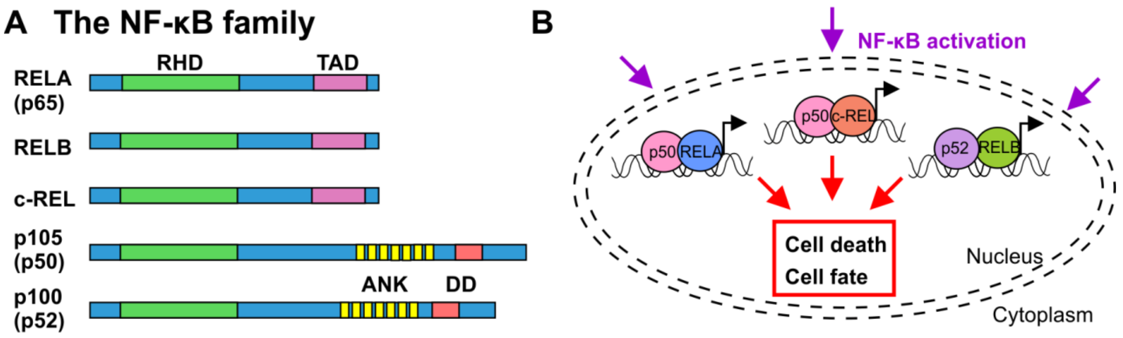


**Figure S1. Schematic view on NF-κB family members and their potential effects on cell fate and survival.** A) Schematic representation of the NF-κB family members. Relevant domains are indicated and alternative nomenclatures are provided in parenthesis. RHD: rel homology domain, TAD: transactivation domain, ANK: ankyrin repeats, DD: death domain. B) Schema showing NF-κB activation triggered during differentiation allowing nuclear translocation of a predominant NF-κB dimer or a particular combination of NF-κB dimers which would induce the gene expression program of a distinct cell fate, or it could result in cell death or in both.

**Figure S2**

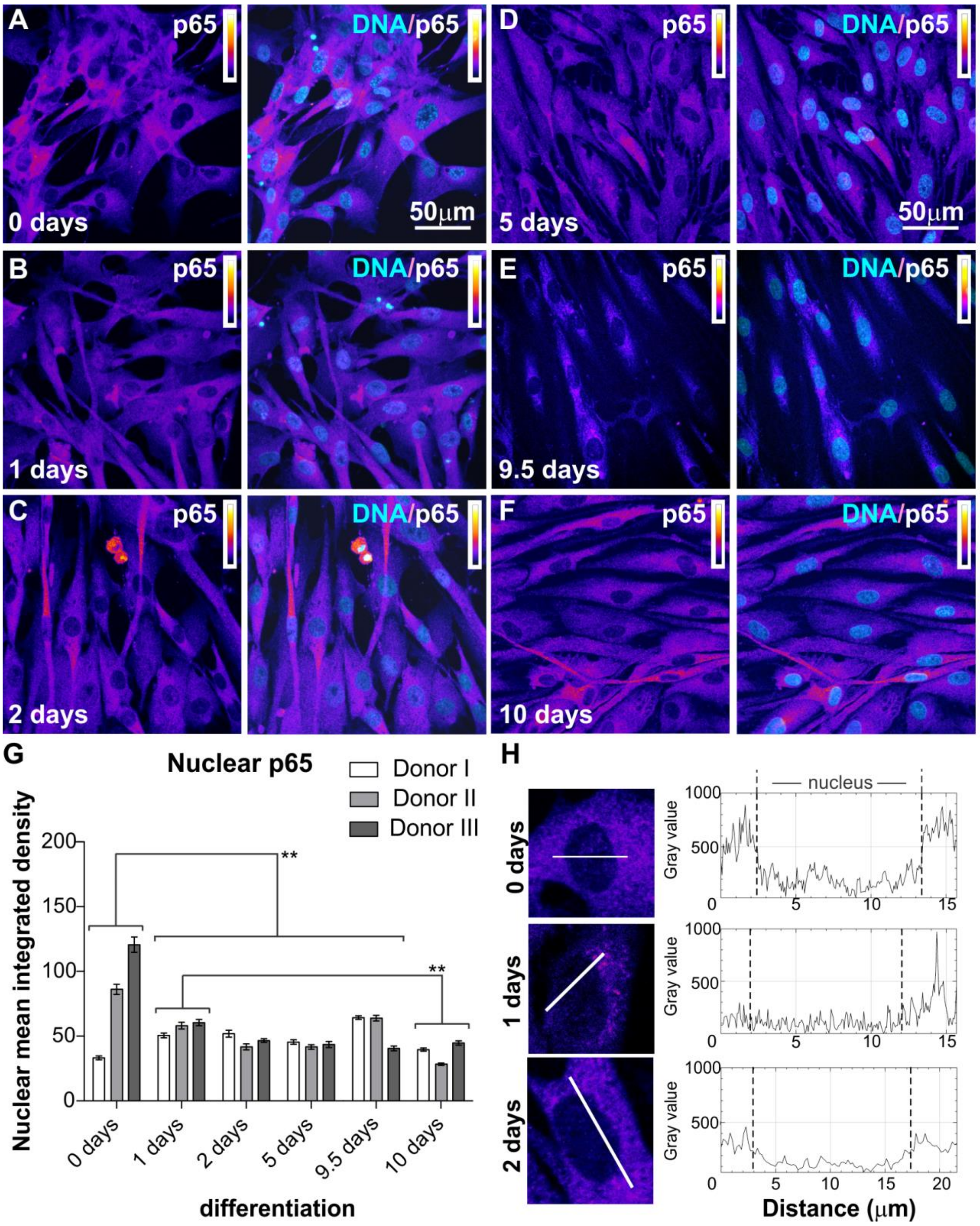


**Figure S2. Immunocytochemical analysis of p65 (RELA).** A-F) NCSC-derived NSCs labeled against RELA after 0, 1, 2, 5, 9.5 and 10 days of glutamatergic differentiation. Each panel shows RELA protein on the left-side and DNA co-localization with DAPI staining on the right-side. Intensity scale indicates white as highest intensity level and black as lowest intensity level. G) Quantification of immunocytochemical analyses showing nuclear mean integrated density of p65 (RELA) during early differentiation (mean ± SEM, n=3). Normality of the data was refuted using Shapiro-Wilk normality test. Non-parametric Kruskal-Wallis (***$p \leq 0.001$) and Bonferroni corrected post-test (**$p < 0.01$) revealed a significant nuclear translocation of NF-κB-p65 at day 0. H) Fluorescence intensity profiles measured at three different time points (0, 1, 2 days of differentiation), for different cells following transects as shown, to reveal the difference between the nuclear and cytoplasmic fluorescence. NCSC: neural crest-derived stem cells, NSCs: neural stem cells, SEM: Standard error of the mean.

**Figure S3**

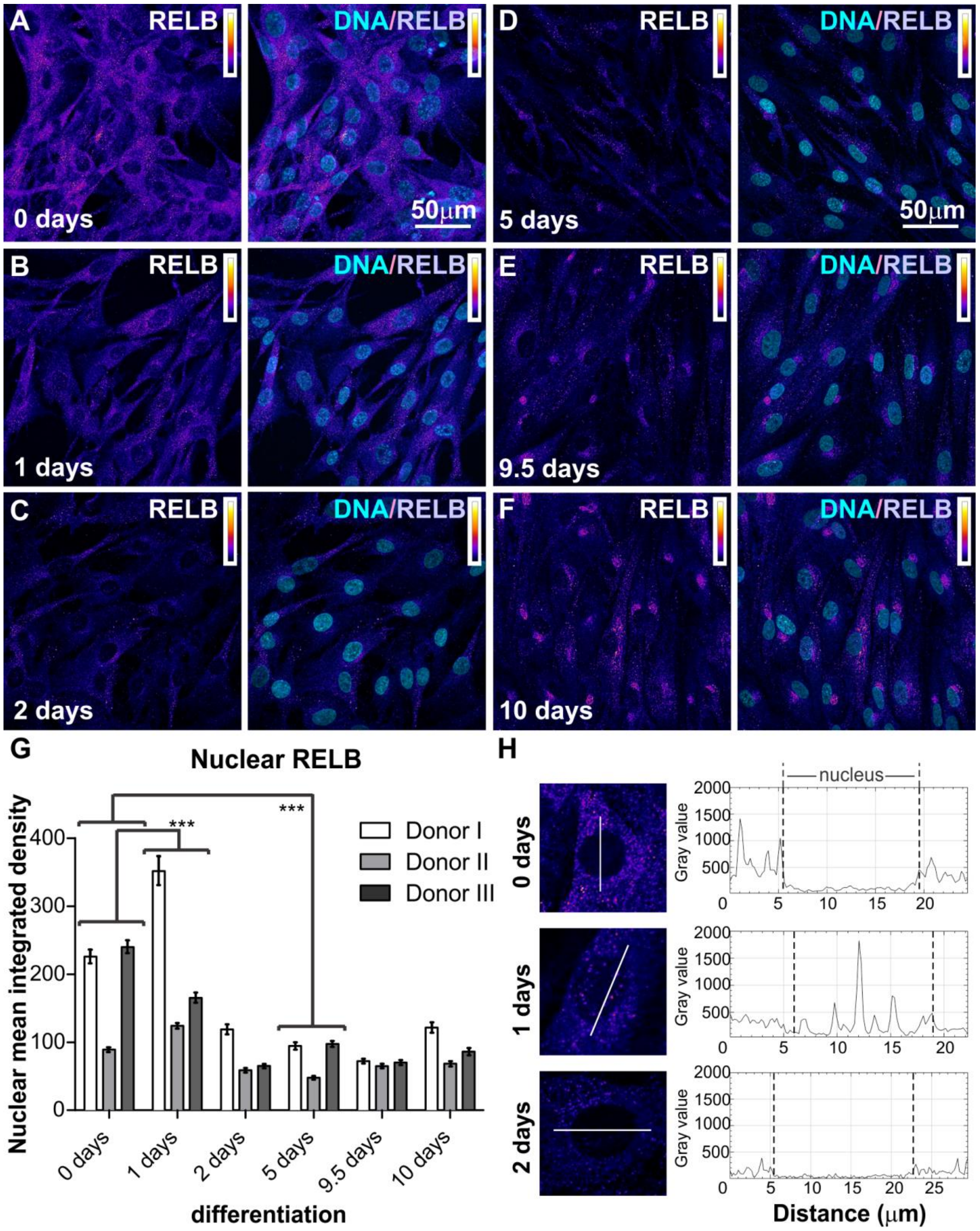


**Figure S3. Immunocytochemical analysis of RELB.** A-F) NCSC-derived NSCs labeled against RELB after 0, 1, 2, 5, 9.5 and 10 days of glutamatergic differentiation. Each panel shows RELB protein on the left-side and DNA co-localization with DAPI staining on the right-side. Intensity scale indicates white as highest and black as lowest intensity levels. G) Quantification of immunocytochemical analyses show nuclear mean integrated density of RELB during early differentiation (mean ± SEM, n=3). Normality of the data was refuted using Shapiro-Wilk normality test. Non-parametric Kruskal-Wallis (***$p \leq 0.001$) and Bonferroni corrected post-test (***$p < 0.001$) revealed a significant peak in nuclear translocation of NF-κB-RELB at day 0. H) Fluorescence intensity profiles measured at three different time points (0, 1 and 2 days of differentiation), for different cells following transects as shown, to elucidate the difference between the nuclear and cytoplasmic fluorescence. NCSC: neural crest-derived stem cells, NSCs: neural stem cells, SEM: Standard error of the mean.

**Figure S4**

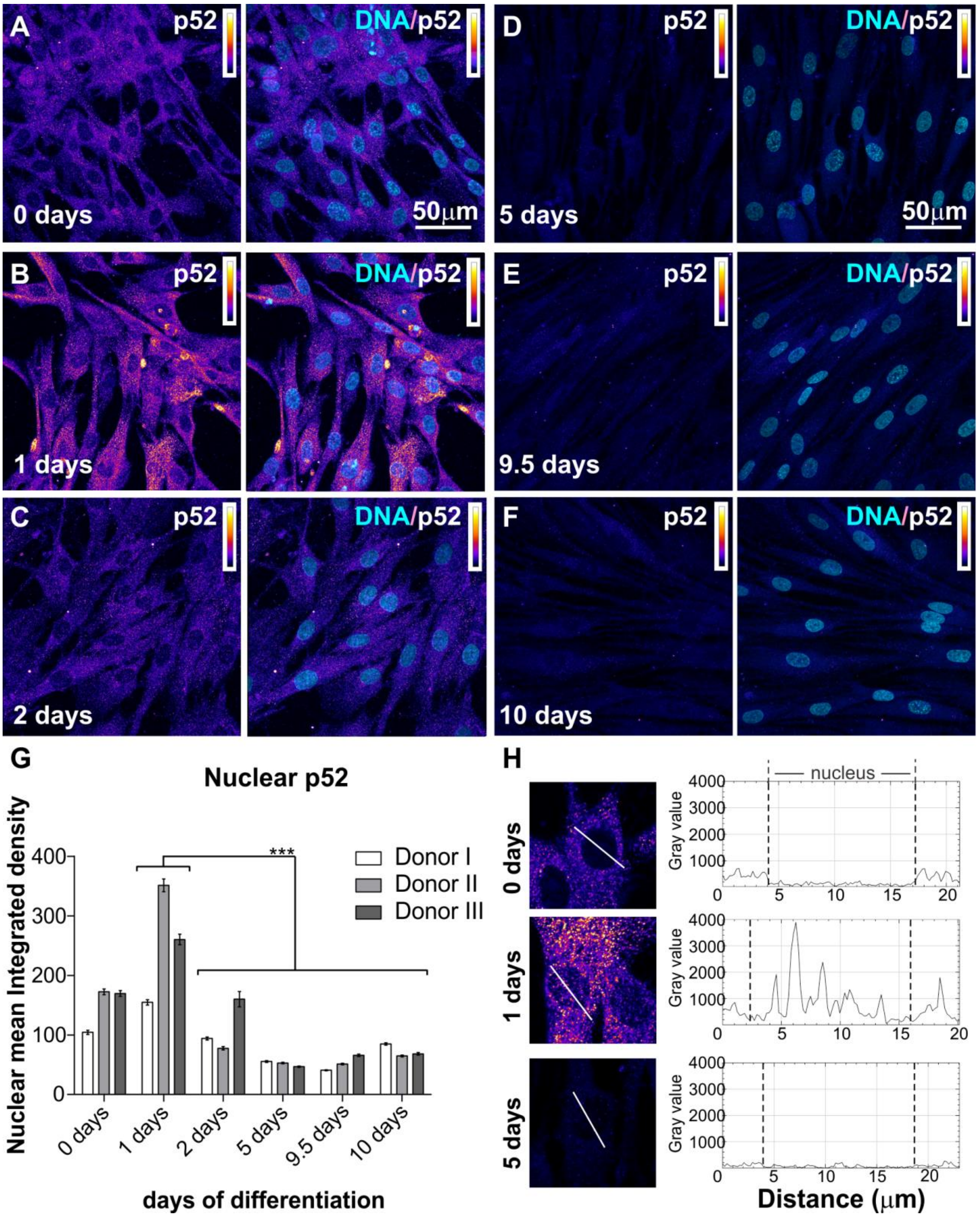


**Figure S4.** Immunocytochemical analysis of p52. A-F) NCSC-derived NSCs labeled against p52 after 0, 1, 2, 5, 9.5 and 10 days of glutamatergic differentiation respectively. Each panel shows p52 subunit on the left-side and the co-localization with DNA on the right-side. Intensity scale indicates white as highest intensity level and black as lowest intensity level. G) Quantification of immunocytochemical analyses showing nuclear mean integrated density of p52 during early differentiation (n=3, mean ± SEM). Normality of the data was refuted using Shapiro-Wilk normality test. Non-parametric Kruskal-Wallis (***$p \leq 0.001$) and Bonferroni corrected post-test (***$p < 0.001$) showed a significant peak of p52 at day 1 significantly different to all later time points (2-10 days). H) Fluorescence intensity profiles measured at different time points (0, 1 and 5 days of differentiation), for different cells following transects as shown, to clearly expose the difference between nuclear and cytoplasmic fluorescence. NCSC: neural crest-derived stem cells, NSCs: neural stem cells, SEM: Standard error of the mean.

**Figure S5**

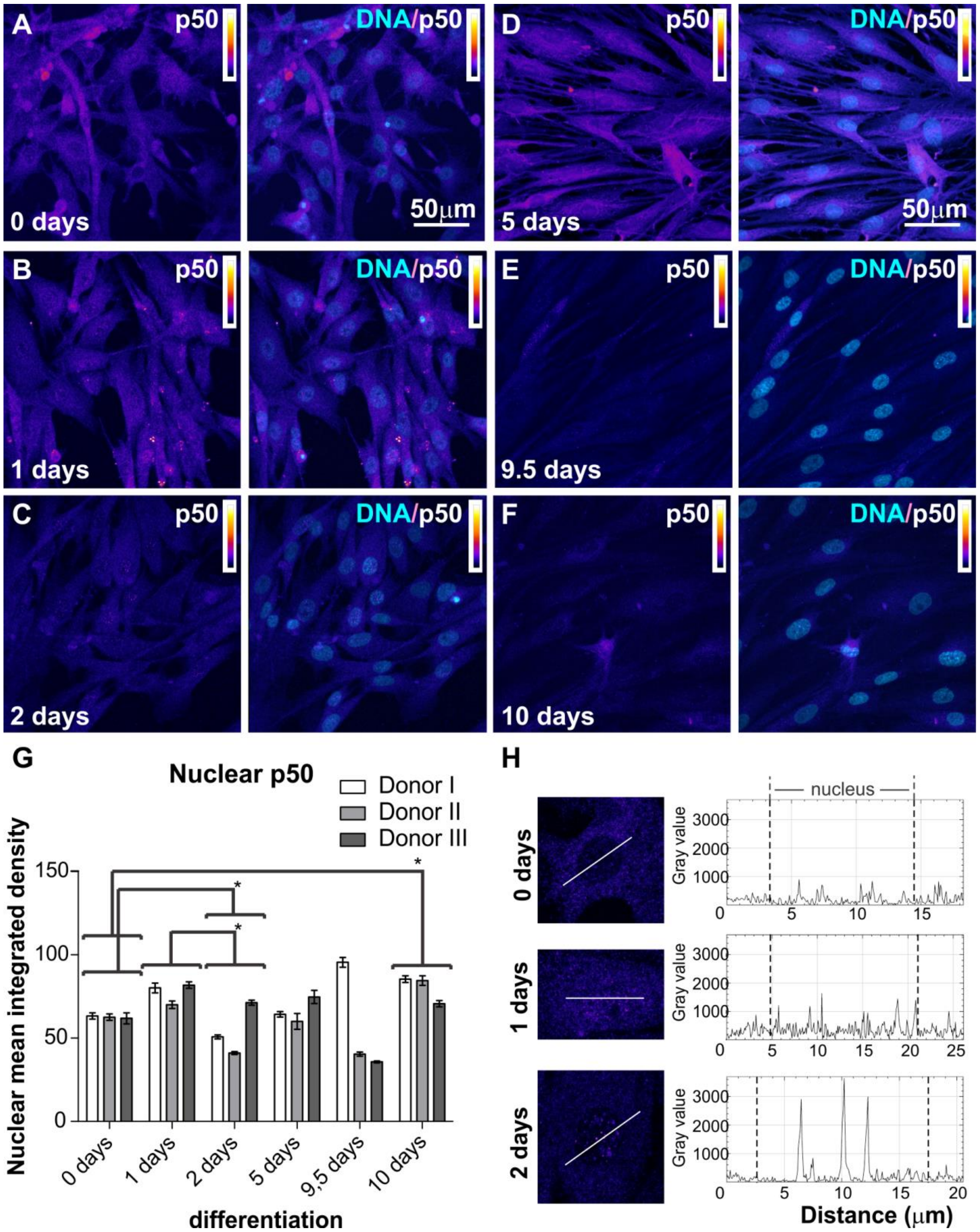


**Figure S5.** Immunocytochemical analysis of p50. A-F) NCSC-derived NSCs labeled against p50 after 0, 1, 2, 5, 9.5 and 10 days of glutamatergic differentiation. Each panel shows p50 protein on the left-side and the co-localization with DNA staining on the right-side. Intensity scale indicates white as highest and black as lowest intensity levels. G) Quantification of immunocytochemical analyses showing nuclear mean integrated density of p50 during early differentiation (n=3, mean ± SEM). Normality of the data was refuted using Shapiro-Wilk normality test. Non-parametric Kruskal-Wallis (***$p \leq 0.001$) and Bonferroni corrected post-test (*$p < 0.05$) showed some significant differences between day 0 and 2 and 10 days, and between day 1 and day 2. H) Fluorescence intensity profiles measured at different time points (0, 1 and 2 days of differentiation), for different cells following transects as shown, to clearly expose the difference between nuclear and cytoplasmic fluorescence. NCSC: neural crest-derived stem cells, NSCs: neural stem cells, SEM: Standard error of the mean.

**Figure S6**

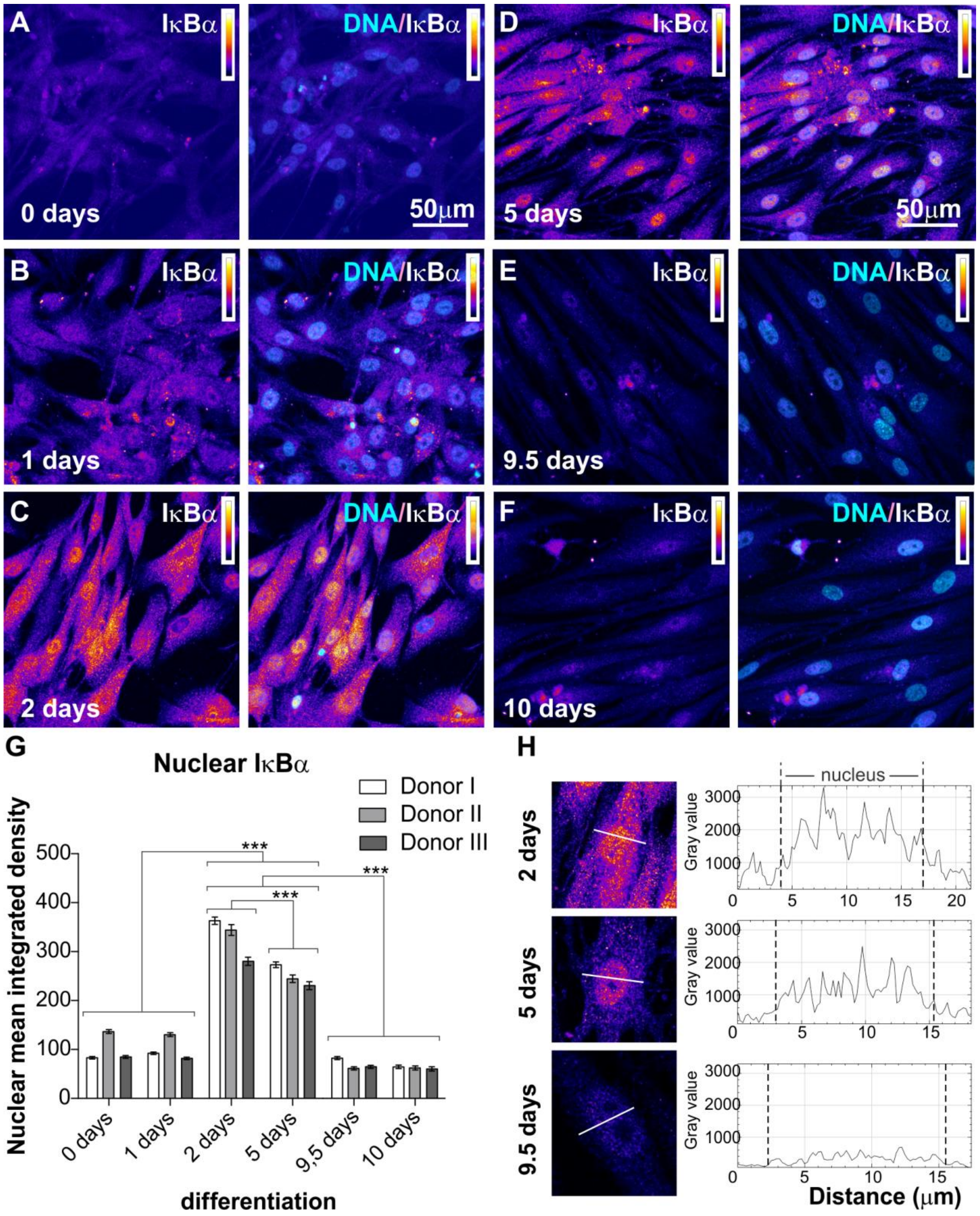


**Figure S6.** Immunocytochemical analysis of IκBα. A-F) NCSC-derived NSCs labeled against IκBα after 0, 1, 2, 5, 9.5 and 10 days of neuronal glutamatergic differentiation respectively. Each panel shows on the left c-REL and co-localization with DNA on the right. Intensity scale indicates white as highest intensity level and black as lowest. G) Quantification of immunocytochemical analyses showing nuclear mean integrated density for IκBα during early differentiation (n=3, mean ± SEM). Normality of the data was refuted using Shapiro-Wilk normality test. Non-parametric Kruskal-Wallis (***$p \leq 0.001$) and Bonferroni corrected post-test (***$p<0.001$) revealed a significantly increased nuclear translocation of IκBα at days 2 and 5. H) Fluorescence intensity profiles measured at three different time points (2, 5 and 9.5 days of differentiation), for different cells following transects as shown, in order to clearly reveal the difference between the nuclear and cytoplasmic fluorescence. NCSC: neural crest-derived stem cells, NSCs: neural stem cells, SEM: Standard error of the mean.

**Figure S7**

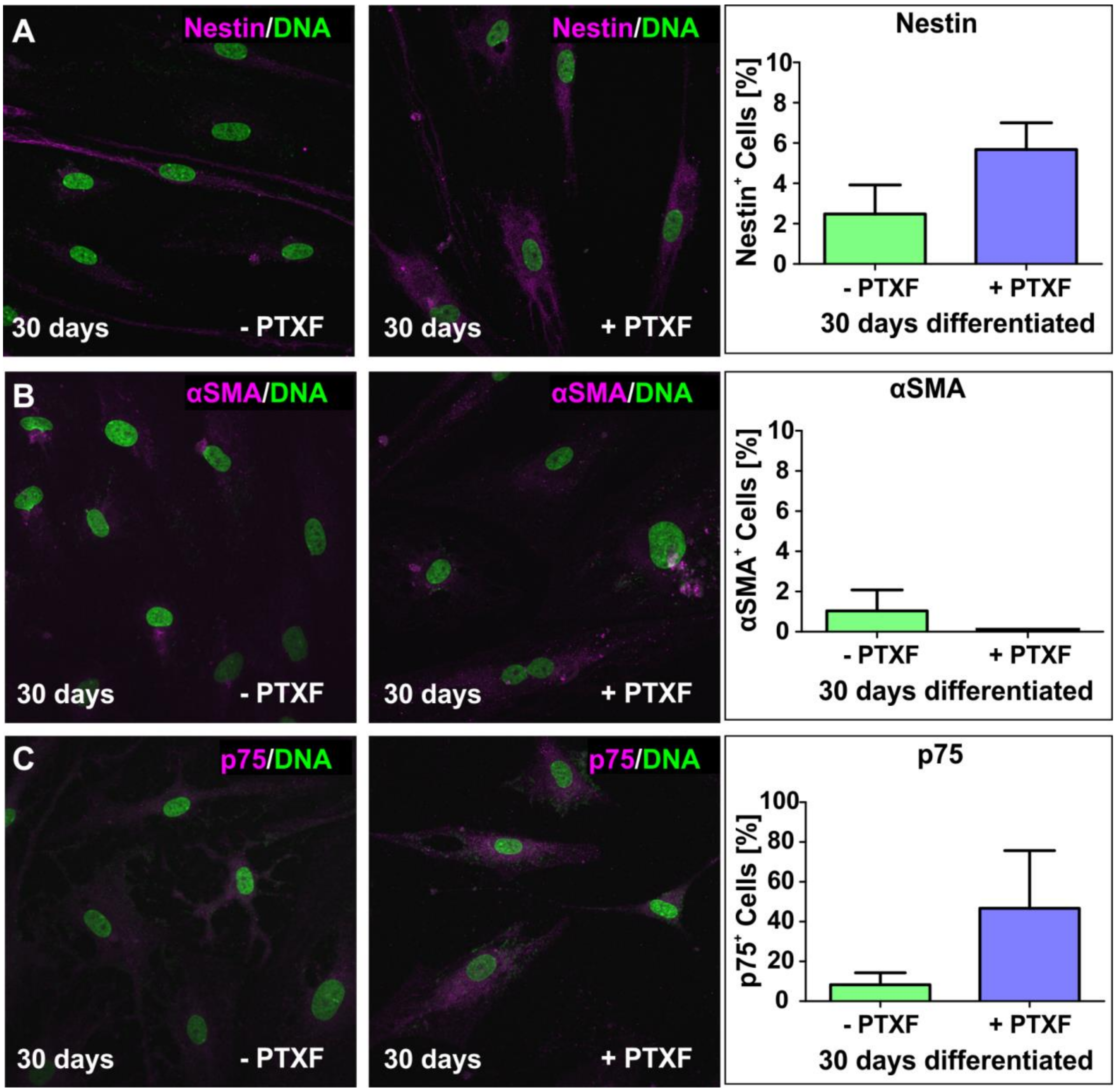


**Figure S7.** Immunocytochemistry assays showing different cell markers after 30 days of glutamatergic neuronal differentiation in the absence or presence of pentoxifylline and respective quantifications. A-B) Neuronally differentiated NCSC-derived NSCs labeled against Nestin, a stemness marker. C) Quantified Nestin$^{+}$ cells shown in percentage suggests a higher tendency of Nestin for the PTXF-treated differentiated-NCSC-derived NSCs (5,68% ± 1,33%), compared to the untreated differentiated NCSC-derived NSCs (2,48% ± 1,44%), however it is not significantly relevant. D-E) Neuronal differentiated NCSC-derived NSCs labeled against αSMA. F) Quantification showing percentage of αSMA$^{+}$ cells which is cero for the differentiated NCSC-derived NSCs treated with PTXF and it is 1,04%(±1,04) in control neurons. No significant difference was observed. G-H) Differentiated NCSC-derived NSCs labeled against p75, a neural crest marker. I) Quantification of the percentage of $p75^{+}$ cells indicates a lower amount of $p75^{+}$ cells (8,25% ± 6,03%), present in the control neurons and a higher tendency of positive cells in the differentiated NCSC-derived NSCs treated with PTXF (46,67% ± 29,06%), however no significant difference. Non-parametric Kruskal-Wallis test ($p<0.10$), no significant differences observed. PTXF: pentoxifylline, NCSC: neural crest-derived stem cells, NSCs: neural stem cells, αSMA: alpha smooth muscle actin.

**Table S1.** Primers sequences for quantitative polymerase chain reaction.

| **Target** | **Primer sequence 5'-3'** |
|---|---|
| Fw- *CSPG4* (chondroitin sulfate proteoglycan type 4*) | CATCCCACTAGAGGCGCAAA |
| Rev- *CSPG4* | CCCAGGAGAGTGGGGAAGTA |
| Fw-*MBP* (Myelin basic protein) | GCGTCACAGAAGAGACCCTC |
| Rev-*MBP* | CTCTGTGCCTTGGGAGGAAG |
| Fw-*PDGFRA* (Platelet derived growth factor receptor alpha) | GAAGAAAACAACAGCGGCCTT |
| Rev-*PDGFRA* | TGTACAACCCTGTGTGGGC |
| Fw-*RPLP0* (Ribosomal Protein Lateral Stalk Subunit P0) | TGGGCAAGAACACCATGATG |
| Rev-*RPLP0* | AGTTTCTCCAGAGCTGGGTTGT |
| Fw-*GAPDH* (Glyceraldehyde-3-phosphate dehydrogenase) | CATGAGAAGTATGACAACAGCCT |
| Rev-*GAPDH* | AGTCCTTCCACGATACCAAAGT |

*also known as transmembrane proteoglycan nerve-glia antigen (NG2).

### Supplementary materials and methods section

#### Extended detailed methods

#### Neuronal differentiation

For neuronal differentiation, NCSCs from three to six donors were expanded and dissociated as described above. Cells were re-suspended in Dulbecco's modified Eagle's medium (DMEM) high glucose (Sigma-Aldrich) containing 2 mM L-glutamine (Sigma-Aldrich), penicillin/streptomycin (1x, Sigma-Aldrich), 10% Fetal Calf Serum (Sigma-Aldrich) and plated at a density of $5x10^4$ cells per 24 well plate followed by cultivation at 37°C, 5% $CO_2$ and atmospheric $O_2$ in a humidified incubator for 2 days. Moreover, 1µM dexamethasone (Sigma-Aldrich), 2 µM insulin (Sigma-Aldrich), 500 µM 3-isobutyl-1-methylxanthine (Sigma-Aldrich), 200 µM indomethacin (Sigma-Aldrich) and 200 µM ethanol were added to the medium to induce neuronal differentiation (neuronal induction medium, NIM) according to (Muller *et al.*, 2015). After 9 days of differentiation cells were induced with 0.5µM retinoic acid (Sigma Aldrich) and 1x N-2 supplement (Gibco, Darmstadt, Germany). Subsequently, the medium was changed by removing half of the volume, followed by addition of fresh pre-warmed NIM containing 1x N-2 supplement (Muller *et al.*, 2015).

#### Immunocytochemistry

Differentiated NCSC-derived NSCs were fixed in phosphate-buffered 4% paraformaldehyde (pH 7.4) for 15 minutes at room temperature (RT) followed by 3 wash steps in phosphate-buffered saline (1xPBS). Cells were permeabilized with 0.02% Triton X-100 and blocked using 5% of appropriate serum or 3% bovine serum albumin for 30 minutes at RT, followed by incubation with primary antibodies for 1 hour at RT. Primary antibodies used were anti-NF-kappa B p65 (1:100, sc-8008, Santa Cruz Biotechnology; 1:200, D14E12, Cell Signaling), anti-c-REL (1:100, sc-70x, Santa Cruz

Biotechnology; 1:400, #4727, Cell Signaling), anti-RELB (1:100, sc-226, Santa Cruz Biotechnology; 1:1600, #10544, Cell Signaling), anti-p50 (1:100, sc-8414, Santa Cruz Biotechnology), anti-p52 (1:100, sc-298, Santa Cruz Biotechnology), anti-IκBα (1:100, sc-371, Santa Cruz Biotechnology), anti-nestin (1:200, MAB5326, Millipore), anti-eurofilament 200 (NF200, 1:200, N4142, Sigma-Aldrich), anti-VGLUT2 (vesicular glutamate transporter 2, 1:200, MAB5504, Millipore), anti-OLIG2 (oligodendrocyte transcription factor 2, 1:250, Q13516, R&D Systems), anti-O4 (1:100, IgM, R&D), Anti-αSMA (alpha smooth muscle actin,1:200, A5691, Sigma), anti-NGFRp75 (nerve growth factor receptor p75, 1:100, sc-6188, Santa Cruz), anti-cleaved caspase-3 (1:300, #9664, Cell Signaling). The secondary fluorochrome-conjugated antibodies were incubated for 1 hour at RT. Secondary antibodies used were goat anti-mouse Alexa 555, goat anti-rabbit Alexa 555, donkey anti goat Alexa 555, and goat anti-mouse-IgM Alexa 555 (1:300, Life Technologies). Nuclear counterstaining was performed with 49,6-diamidino-2-phenylindole (DAPI; 1 μg/ml; Sigma-Aldrich) for 15 min at RT. Fluorescence imaging was performed using a confocal laser scanning microscopy (LSM 780; Carl Zeiss, Jena, Germany) and analyzed using ZEN software from the same provider or ImageJ.
